# Cholesterol interaction sites on the transmembrane domain of the hedgehog signal transducer and Class F G protein-coupled receptor Smoothened

**DOI:** 10.1101/383539

**Authors:** George Hedger, Heidi Koldsø, Matthieu Chavent, Christian Siebold, Rajat Rohatgi, Mark S. P. Sansom

## Abstract

Transduction of hedgehog signals across the plasma membrane is a key process during animal development. This is facilitated by the Class F G-protein-coupled-receptor (GPCR) Smoothened (SMO), a major drug target in the treatment of basal cell carcinomas. Recent studies have suggested that SMO is modulated *via* interactions of its transmembrane (TM) domain with cholesterol. Long time scale (>0.35 ms of simulation time) molecular dynamics simulations of SMO embedded in two different cholesterol containing lipid bilayers reveal direct interactions of cholesterol with the transmembrane domain at regions distinct from those observed in Class A GPCRs. In particular the extracellular tips of helices TM2 and TM3 form a well-defined cholesterol interaction site, robust to changes in membrane composition and in force field parameters. Potential of mean force calculations for cholesterol interactions yield a free energy landscape for cholesterol binding. Combined with analysis of equilibrium cholesterol occupancy these results reveal the existence of a dynamic ‘greasy patch’ interaction with the TM domain of SMO, which may be compared to previously identified lipid interaction sites on other membrane proteins. These predictions provide molecular level insights into cholesterol interactions with a biomedically relevant Class F GPCR, suggesting potential druggable sites.

## Introduction

The Smoothened (SMO) receptor is a critical component of the hedgehog signaling cascade, which controls a variety of key developmental processes, including human embryonic tissue patterning and regulation of adult stem cells (Briscoe and Thérond, 2013). Aberrant activation of SMO causes uncontrolled hedgehog signaling and a variety of cancers (Wu et al., 2017). As such SMO is of major academic and pharmaceutical interest, and is the target of two FDA approved drugs for treating basal cell carcinomas, vismodegib (Dlugosz et al., 2012) and sonidegib (Burness, 2015).

SMO is a member of the Frizzled-class of G-protein coupled receptors (GPCRs) and is found at the plasma membrane. Its structural architecture consists of an extracellular cysteine rich domain (CRD), stacked on top of a short linker domain (LD), and a hepta-helical transmembrane domain (7TMD) (Byrne et al., 2016; Zhang et al., 2017), (Figure 1). To date, eleven crystal structures containing the SMO 7TMD have been solved (Byrne et al., 2018), revealing structural similarity to the presumed inactive state of Class A GPCRs (Wang et al., 2013). However, SMO exhibits < 10% sequence identity to Class A GPCRs, and is missing the canonical D[E]RY and NPxxY motifs implicated in the signaling mechanisms of Class A receptors (Wang et al., 2013).

**Figure 1.**
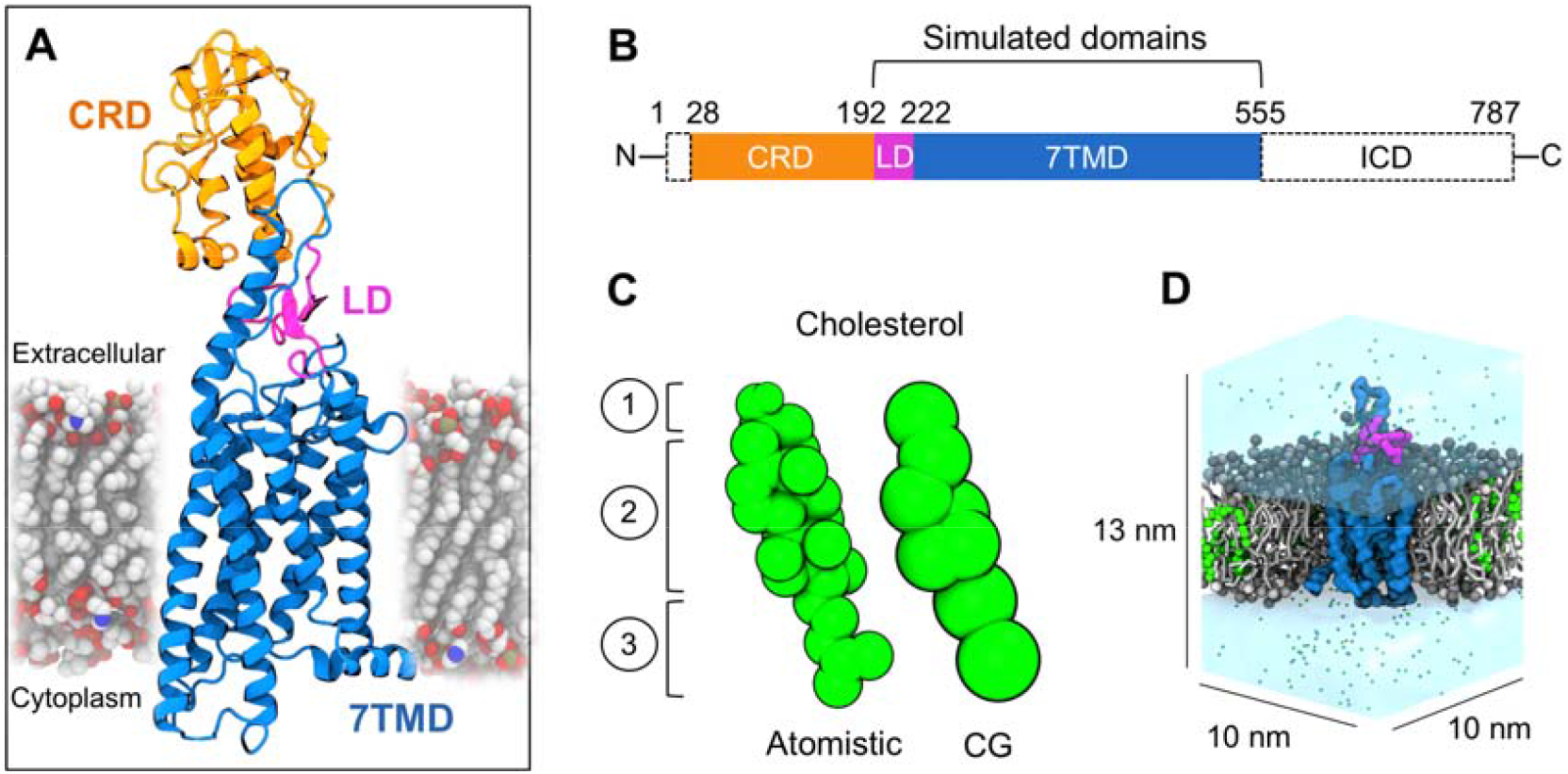
System overview. (A, B) Molecular architecture (PDB: 5L7D) of the multi-domain human SMO protein, consisting of an extracellular CRD (orange), Linker domain (magenta), 7TM domain (blue), and structurally unresolved intracellular domain (ICD). (C) Sphere representation of atomistic and CG Martini cholesterol, depicting the polar hydroxyl head group (1), hydrophobic cyclic ring system (2), and iso-octanyl tail (3). (D) Cross-section through a CG system showing the simulated SMO construct (Linker domain + 7TMD) embedded in a cholesterol (green) containing phosphatidylcholine (PC) lipid bilayer. Water is depicted as a transparent surface, and ions shown as sphere representations.

While the mechanism of SMO activation in the course of physiological Hh signaling has been a long-standing mystery, a number of recent studies have suggested that cellular cholesterol plays a central role in SMO activation and transduction of the Hh signal across the membrane (Blassberg et al., 2016; Byrne et al., 2018; Cooper et al., 2003; Huang et al.; Luchetti et al., 2016; Myers et al., 2013). Reduced cellular cholesterol levels, as seen in Smith-Lemli-Opitz syndrome (SLOS), or as produced when cells are treated with methyl-β-cyclodextrin (MβCD), lead to decreased SMO activity and blunted Hh responses in target cells (Blassberg et al., 2016; Cooper et al., 2003). In addition, interesting synergistic effects on SMO function have been reported (Gordon et al., 2018) between statins, used to suppress cholesterol biosynthesis, and the SMO antagonist vismodegib, in the treatment of Medulloblastoma. Further, oxysterols, hydroxylated metabolites of cholesterol, were shown to function as direct SMO agonists, suggesting SMO could function as a sterol receptor (Nachtergaele et al., 2012).

The near full-length crystal structure of the protein revealed an extracellular cholesterol binding site located within the CRD (Byrne et al., 2016). Binding of cholesterol at this site was subsequently shown to be both necessary and sufficient for activation of SMO and hedgehog signaling (Huang et al.; Luchetti et al., 2016). However, constitutively active truncations of SMO entirely lacking the CRD and SMO mutants that cannot bind sterols through the CRD (Blassberg et al., 2016; Briscoe and Thérond, 2013) still depend on the presence of membrane cholesterol for their activity (Myers et al., 2017), suggesting that cholesterol may also regulate SMO activity through a second site within the transmembrane domain. However at present no molecular level detail exists on the possible location of such an interaction.

Coarse-grained (CG) (Marrink et al., 2007) molecular dynamics simulations (Figure 1) enable the study of lipid interactions with integral membrane proteins (Hedger et al., 2016b). This approach provides access to the long simulation timescales and large system sizes required for thorough sampling of lipid-protein interaction space (Corradi et al., 2018). For example, this approach has previously been applied to predict the location of PIP2 binding sites on Kir channels (Stansfeld et al., 2009), supported by a subsequently determined PIP_2_-bound crystal structure (Hansen et al., 2011), and to characterize the structural, dynamic, and energetic aspects of cardiolipin interactions with the ADP/ATP translocase (Duncan et al., 2018; Hedger et al., 2016b), in agreement with available NMR and crystallographic data. More recently, the predicted locations of PIP2 binding sites on Class A GPCRs have been shown to agree with the results of site directed mutagenesis and native mass spectrometry experiments (Yen et al., 2018). A number of other studies have compared the accuracy of the CG simulation approach to available experimental data on lipid binding for a variety of membrane proteins (Arnarez et al., 2013a; Arnarez et al., 2013b). In the case of cholesterol, the model recently identified both novel and established cholesterol binding sites on the Adenosine A2A receptor (Rouviere et al., 2017), as well on as the dopamine transporter (Zeppelin et al., 2018). For a comprehensive review of cholesterol interactions studied by molecular dynamics simulations see (Grouleff et al., 2015), and for a comparative analysis of cholesterol interactions with a range of membrane proteins see (Lee, 2018).

Here we present multiscale simulations of human SMO embedded in a cholesterol-containing membrane environment (Figure 1D…Table 1.) Cholesterol interactions are addressed *via* equilibrium CG simulations in both a simple two-component bilayer, and in a more complex *in vivo* mimetic membrane. We also calculate potentials of mean force to estimate the free energy of the SMO-cholesterol interaction in a bilayer environment. Our data predict a direct interaction of cholesterol with the transmembrane domain of SMO, at defined locations.

## Results

All simulations were conducted using the GROMACS 4.6.5 (http://www.gromacs.org) simulation package (Hess et al., 2008). An overview of simulations performed is detailed in Table 1.

**Table 1.** Overview of the simulations performed. The box size for equilibrium simulations was 10 × 10 × 13 nm^3^. The tail saturation pattern for phospholipids was 1-palmitoyl-2-oleoyl.

| Description | Membrane composition | Duration |
| --- | --- | --- |
| 7TMD and linker domain, CG | PC (80%) + Chol (20%) | 8 x 10 μs |
| 7TMD and linker domain, CG, virtual site cholesterol (Melo et al., 2015) | PC (80%) + Chol (20%) | 3 x 10 μs |
| Alternative conformation (Huang et al.) of the 7TMD and linker domain, CG | PC (80%) + Chol (20%) | 8 x 10 μs |
| 7TMD and linker domain, CG | PC (80%) + Chol (20%) | 1 x 100 μs |
| SMO 7TM and linker domain, AT | PC (80%) + Chol (20%) | 3 x 0.2 μs |
| 7TM PMF calculations, CG | PC (100%) | 32 x 1 μs<br>per PMF |
| 7TMD and linker domain, CG | Outer leaflet: PC (60%) + PE (15%) + Chol (25%)<br>Inner leaflet: PC (10%) + PE (40%) + PS (15%) + PIP<sub>2</sub> (10%) + Chol (25%) | 8 x 10 μs |

*The box size for equilibrium simulations was 10 x 10 x 13 nm<sup>3</sup>. The tail saturation pattern for phospholipids was 1-palmitoyl-2-oleoyl.*

### Cholesterol interacts directly with the SMO transmembrane domain at defined regions

The CG model of the 7TMD+LD of SMO were initially embedded in a PC:Chol (80%/20%) membrane (Table 1), with the cholesterol content chosen based on mammalian lipidomics estimates (Ingolfsson et al., 2014; van Meer et al., 2008). Eight separate systems with different initial lipid configurations were constructed, and used to initiate 8 × 10 μ of coarse-grained molecular dynamics (CGMD) simulations.

In all cases cholesterol approached the protein on a sub-microsecond timescale, forming direct interactions with the transmembrane domain. The average molar composition of the first lipid interaction shell was 7.8 cholesterol and 26.7 PC molecules (22% cholesterol content) over the simulated time course (Figure S2). This indicates no significant annular enrichment in cholesterol compared to the bulk membrane (20% cholesterol content). Rather than form uniform interactions across the membrane exposed surface in a non-specific fashion, cholesterol molecules were seen to localize at defined regions around the protein. Calculation of the time-averaged probability density for cholesterol around the protein revealed a particularly intriguing propensity for interaction with TM2/3 (Figure 2A). The extracellular portions of these helices (TM2/3e) together with ECL1 form a groove-like architecture on the protein surface into which a single cholesterol molecule slots (Figure 2B). Consistently high density at this site was observed across all eight simulation site, independent of the starting configurations (Figure S3). This contrasts with the more diffuse probability densities observed at other regions.

In order to assess the robustness of our observations of SMO-cholesterol interaction patterns in a simple two-component bilayer to changes in lipid composition, we repeated the initial 8 × 10 β set of simulations in a five-component lipid bilayer containing PC, PE, PS, PIP2, and cholesterol. These lipids, their distribution between leaflets, and their relative percentages were chosen to mimic a simplified *in vivo* plasma membrane composition (Table 1). No significant differences in cholesterol interactions were seen compared to the initial two-component (PC + cholesterol) membranes (Figure 2). This demonstrates that our simulation procedure reproducibly identifies a pattern of cholesterol interactions with SMO, especially the TM2/3e site, in two independent extensive ensembles of simulations with different lipid bilayer compositions.

**Figure 2.**
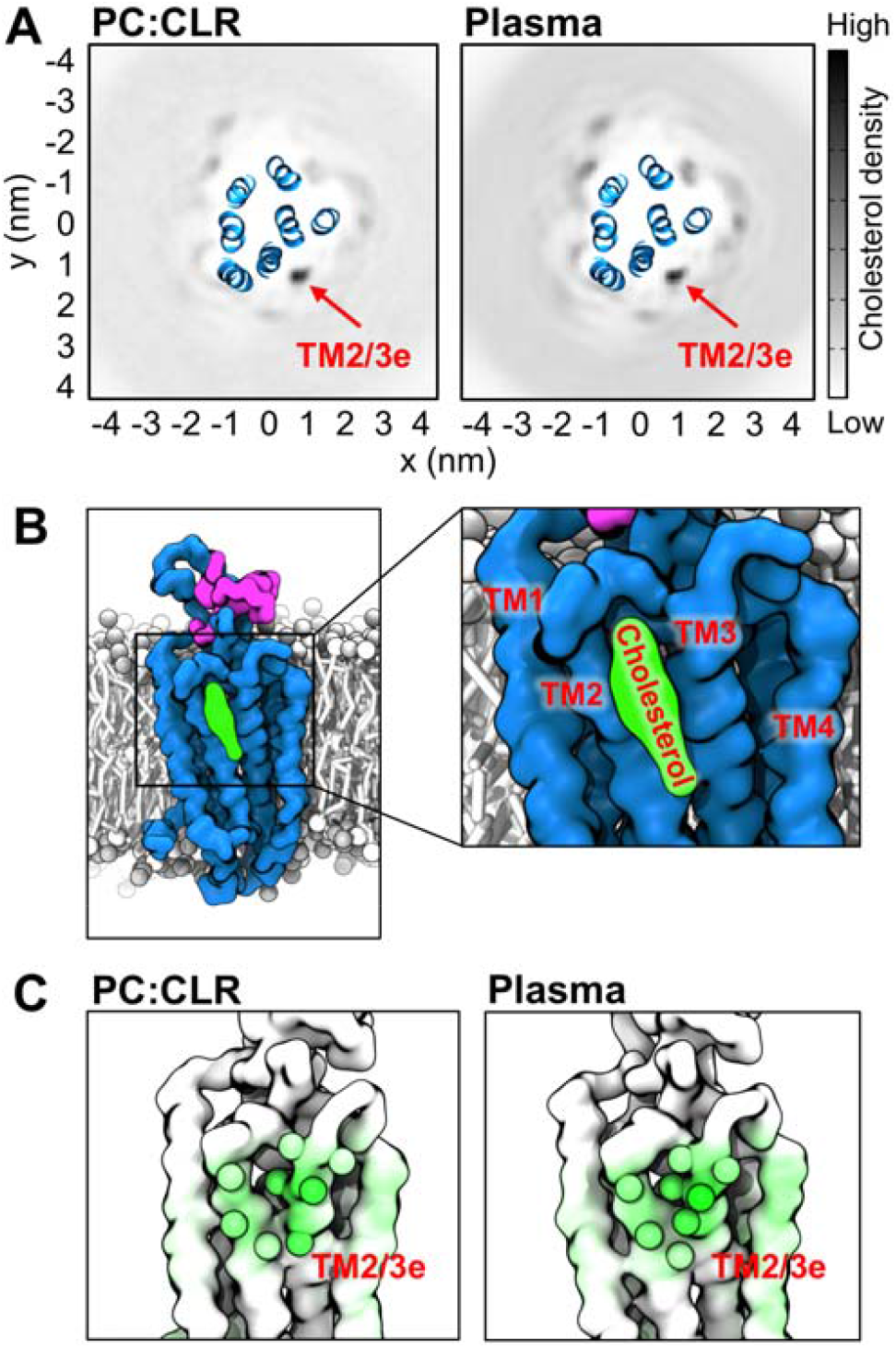
Cholesterol forms direct interactions with SMO and localizes to defined regions. (A) 2D time-averaged density projections for membrane cholesterol around SMO in a simple two component membrane composition (left) and a plasma membrane mimetic (right). (B) Final snapshot (t = 10 μs) from a simulation showing cholesterol bound at the TM2/3e site. (C) View onto the TM2/3e site with simulation snapshots of the protein colored from white (no contact) to green (high contact) according to the degree of interaction with cholesterol. Data for both membrane compositions were averaged over 8 independent CGMD simulations each of 10 μs duration, initiated from different random lipid configurations. See Table 1. for further details of the membrane compositions employed.

To assess the molecular interactions giving rise to this density, the number of contacts formed between cholesterol and each residue of the protein were calculated over the simulated time course. Mapping these contacts onto the protein structure revealed cholesterol interaction hotspots on the membrane-exposed surface (Figure 3; Figure S4). Within the TM2/3e site, the highest degree of cholesterol contact was formed with V276, I279, A283, M286, L312, S313, I316, I317, and I320 (Figure 3B). The majority of these contacts occurred with the hydrophobic moieties of cholesterol, whilst a degree of interaction was also seen for the headgroup hydroxyl with S313. The triad of isoleucine residues formed particularly high levels of interaction, a trend which has been observed for cholesterol binding sites on other membrane proteins (Gimpl, 2016). Interestingly, although well-defined density was absent around TM4, the contact analysis coupled with visual inspection of the trajectories showed a moderate level of more dynamic interaction within this region. We explored the robustness of these predicted contacts by also using an alternative set of CG cholesterol parameters employing virtual sites (Melo et al., 2015), performing three independent replicates each of 10 μs duration. The same residue-by-residue cholesterol interaction pattern was observed compared to the standard parameter set (de Jong et al., 2013a; Marrink et al., 2008) (Figure S5).

**Figure 3.**
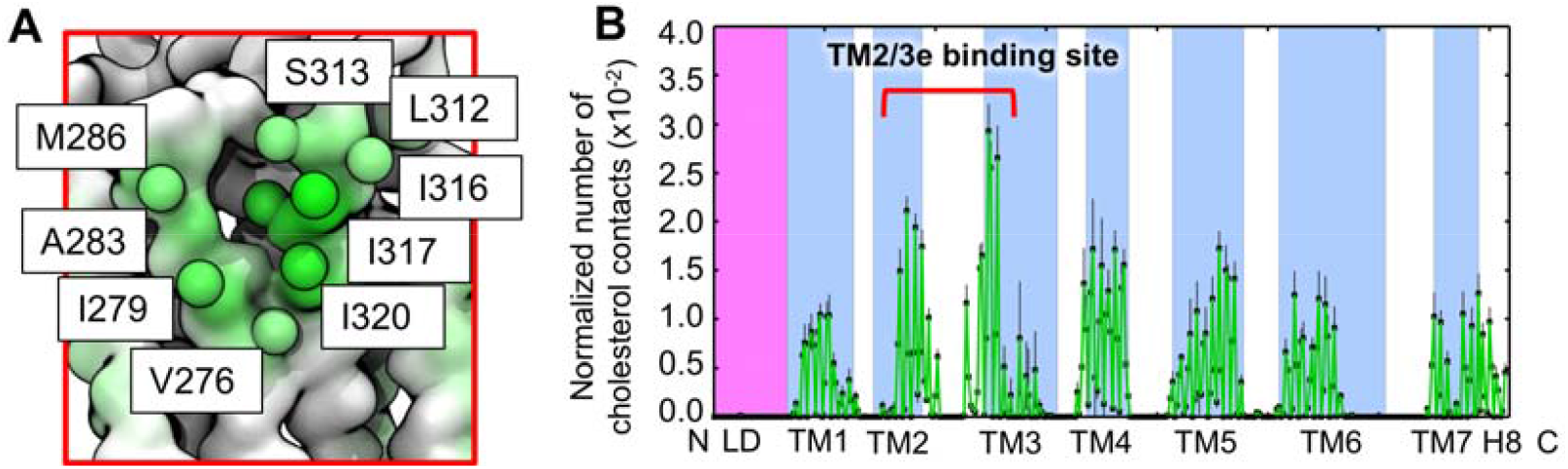
Per residue time-averaged cholesterol contacts with SMO. (A) Zoom-in on the putative TM2/3e cholesterol binding pocket with binding site residues labeled and depicted as spheres, colored from white (no contact) to green (high contact) according to the degree of cholesterol interaction. (B) Normalized number of cholesterol contacts for each residue of the protein, with the standard deviation (n=8) denoted by black error bars. The linker domain and transmembrane helices are delineated by magenta and blue boxes. Contacts were calculated using a 6 Å distance cutoff to define ‘contact’, based on the radial distribution function for CG Martini lipid-protein interactions.

Recently two additional structures of *Xenopus laevis* SMO emerged, bound to cholesterol and to cyclopamine at the CRD (Huang et al.). Both structures exhibited an alternative 7TMD conformation in which the intracellular portion of TM6 adopts an arrangement in which the intracellular tip moves outward by several Ångstrom compared to previous SMO structures (Byrne et al., 2016; Wang et al., 2014; Wang et al., 2013; Weierstall et al., 2014; Zhang et al., 2017). We subjected the 7TMD of the cyclopamine bound structure (PDB id: 6D32) to 8 × 10 μs of CGMD, performed in the same manner already described. The same TM2/3e binding site as seen in the human SMO structure was also observed in the alternative *X. laevis* structure (Figure S6). Interactions at the TM5/6 region were similar in both sets of simulations, exhibiting some degree of more transient interaction, but no stable localization. Notably, we did not observe spontaneous entry of cholesterol into the core of the protein between TM5/6 as has been proposed (Huang et al.).

### Long time scale simulation of cholesterol dynamics

To enable assessment of the dynamics of the interaction and better test system convergence, we extended one CG simulation from 10 s to 100 s. No significant evolution in cholesterol interaction patterns were observed on this long timescale compared to the ensemble of shorter simulations (Figure S7), again indicating convergence of the system properties under consideration.

Calculation of the time-dependent occupancy of the TM2/3e site revealed the site remained occupied by a cholesterol molecule for close to the entire duration of the 100 μs simulation (Figure 4A). This high level of occupancy could result either from extended binding of a single cholesterol molecule, or from rapid exchange events between multiple different molecules. We assessed this question by decomposing the occupancy data to form an interaction matrix for individual cholesterol molecules, showing the occupancy between each individual cholesterol index in the simulation and the TM2/3e site (Figure 4B). The dynamic nature of the interaction is apparent, with exchange between different cholesterol molecules at the TM2/3e site on a sub-microsecond timescale (Figure 4C) (Supplementary Movie S1) leading to exhaustive sampling by all 54 molecules over the course of the 0.1 ms simulation. Binding events ranged from transient interactions in the order of tens of nanoseconds, through to extended interactions of 1 μs or more. Visual inspection of the longest binding event (7 μs duration) revealed that even within extended interaction events, the bound cholesterol molecule was dynamically localized and frequently rotated around its long axis, alternately exposing its rough β-face to the membrane and binding site. This observation of a dynamic interaction with a ‘greasy patch’, as opposed to ‘rigid binding’ concurs with the findings of Lyman and colleagues for the Adenosine A2A receptor (Rouviere et al., 2017), and a range of other cholesterol binding membrane proteins (Grouleff et al., 2015), and may in turn correlate with the absence of well-defined cholesterol density at this region in available crystal structures of SMO (Byrne et al., 2016; Wang et al., 2013).

**Figure 4.**
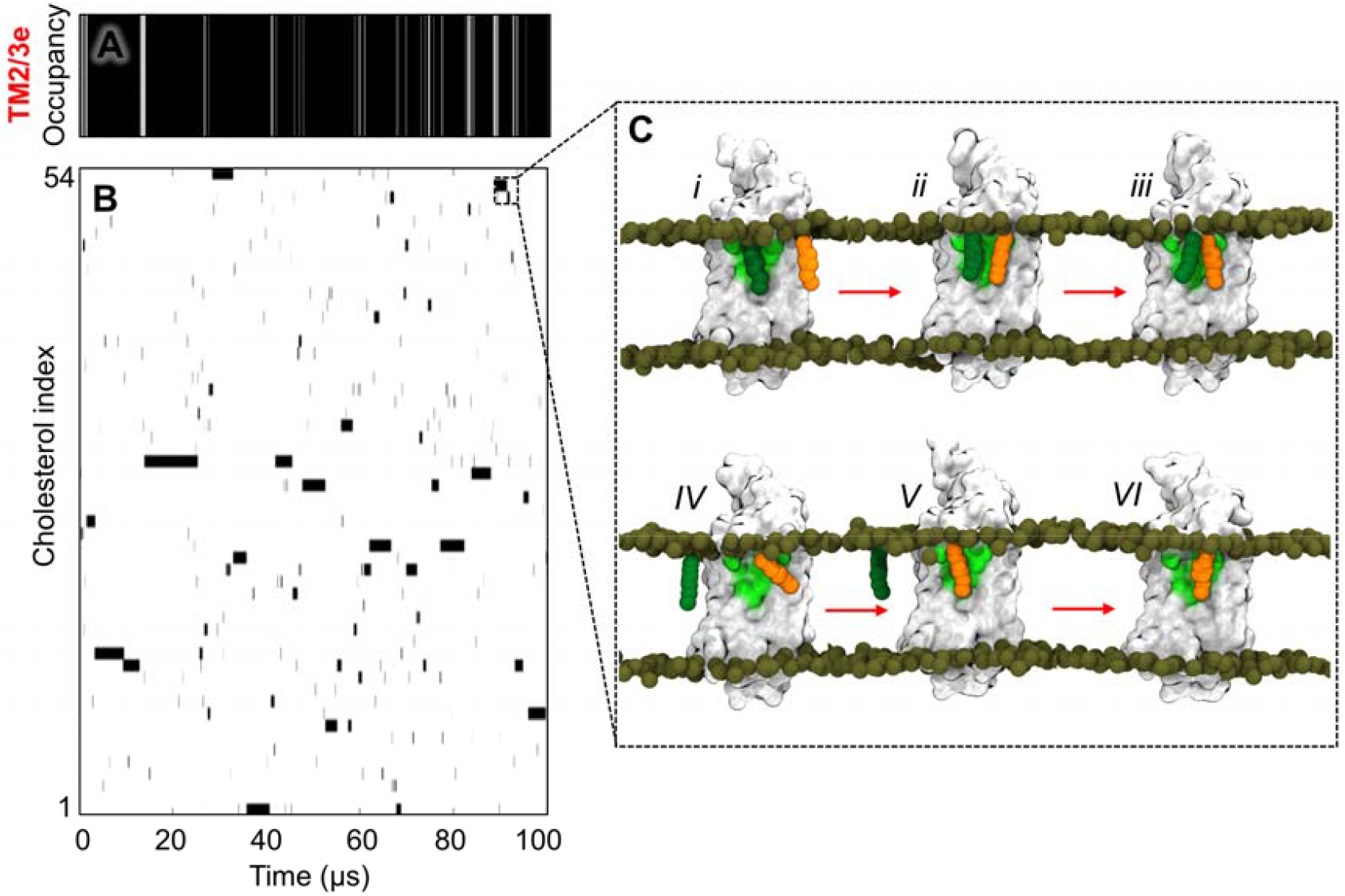
Dynamics of cholesterol interaction within long time-scale MD. (A) Occupancy of the TM2/3e site over time, with black=occupied, white=unoccupied. (B) Interaction matrix showing the occupancy data for each individual cholesterol molecule with TM2/3e. This data shows that the TM2/3e site is occupied for almost the entire duration of the 100 μs simulation, and that this is due to multiple rapid exchanges between different cholesterol molecules rather than a single long-timescale binding of one cholesterol. (C) A series of snapshots are shown (right) depicting an exchange event between two cholesterol molecules (orange, dark green). Occupancy data were calculated as previously described by us (Chavent et al., 2018) using a distance cutoff of 8 Å between the centre-of-mass of the TM2/3e site and the centre-of-mass of each cholesterol molecule, in keeping with (Arnarez et al., 2013b; Hedger et al., 2016a).

### All-atom simulations also demonstrate a direct cholesterol interaction and reveal additional molecular details

To better investigate the atomic level details of the interaction, three snapshots from the CG simulations with cholesterol bound at TM2/3e were converted to atomic resolution. Each simulation was equilibrated for 10 ns with position restraints on the protein, before being run for 200 ns of unbiased molecular dynamics. The protein remained stable with Cα root mean square deviation (R.M.S.D.) of the TM helices plateauing at ~ 0.2 nm for all three repeats (Figure S8). The predominant structural fluctuations were seen in disordered loop regions, and particularly the long ECL3 connecting TM6 and TM7. These observations agree well with our previous atomistic simulations of the full-length SMO construct (Byrne et al., 2016). In all three simulations the cholesterol molecule at site TM2/3e remained bound, undergoing frequent rotation about its long-axis, alternately exposing the methyl groups of its rough β-face to the membrane and binding site, as observed in the CG simulations (Figure 5A) (Supplementary Movie S2). In two cases the cholesterol molecule underwent partial dissociation (Figure 5B: red arrows) before re-binding to adopt is previous orientation, emphasizing the dynamic nature of the interaction. Interestingly, E292, not identified in the CG simulations, was also seen to form interactions with the hydroxyl group of the bound cholesterol, and the adjacent S313 (Figure 5C). This additional observation underscores the value in adopting a serial multiscale approach (Stansfeld and Sansom, 2011) in the investigation of lipid-protein interactions.

**Figure 5.**
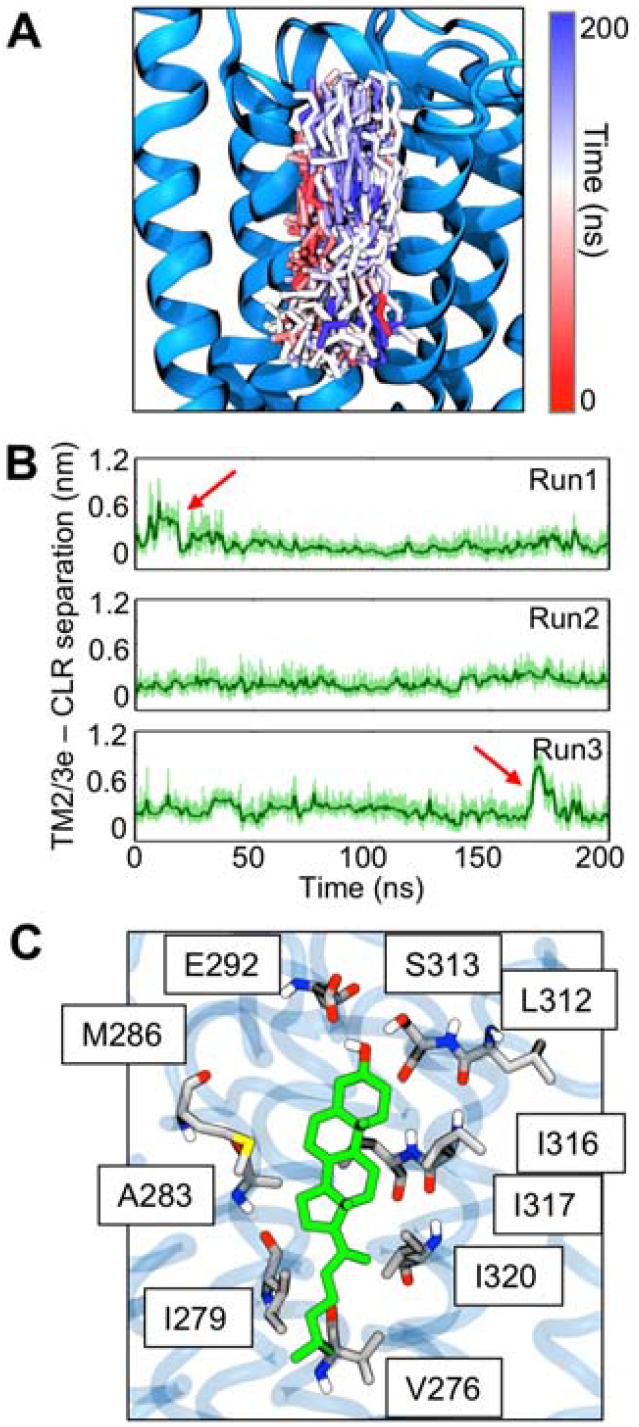
Cholesterol interactions within all-atom simulations. (A) Snapshot of SMO embedded in a lipid bilayer, showing 600 evenly distributed structures of cholesterol (stick representation) at the TM2/3e binding pocket during 3 × 200 ns of atomistic simulation. Structures are shown colored according to simulation time. (B) Distance between the centre-of-mass of cholesterol and the TM2/3e binding pocket. (C) Final simulation snapshot (t = 200 ns) showing the arrangement of key binding site residues.

### Potential of mean force calculations for lateral cholesterol interaction

To assess the strength and selectivity of cholesterol interaction at TM2/3e we calculated the potential of mean force (PMF) for the lateral interaction of membrane cholesterol with the site. The PMF describes the change in free energy between two species along a particular reaction coordinate, and is derived from the probability distribution along this coordinate (Roux, 1995). A steered MD simulation was performed in which a force was applied to pull the bound cholesterol molecule away from its binding site into the bulk membrane. This generated a lateral 1D reaction coordinate (r) perpendicular to the protein surface, ranging from the bound to unbound state. Umbrella sampling was then applied to calculate the free energy profile along this coordinate, with the reaction coordinate *r* defined as the distance between the centre-of-mass of the TM2/3e binding site and the cholesterol molecule.

The profile uncovered a maximal well-depth of ca. −10 kJ/mol at TM2/3e, for both the standard (Marrink et al., 2008) and virtual site (Melo et al., 2015) cholesterol parameters (Figure 6). In contrast repeating the calculation for a separate site on the intracellular portion of the protein (the CCM, which has been suggested to bind cholesterol in Class A GPCRs (Hanson et al., 2008)) which is not predicted to bind cholesterol from our equilibrium simulations, yielded a well-depth < 2.5 kJ/mol (RT), indicating no significant interaction at this site (Figure S9). Repeating the same calculation at TM2/3e for PC, PE, and PS lipids, which comprise a significant portion of plasma membrane lipids (van Meer et al., 2008), likewise showed no significant interaction, demonstrating selectivity of the site for cholesterol (Figure 6).

**Figure 6.**
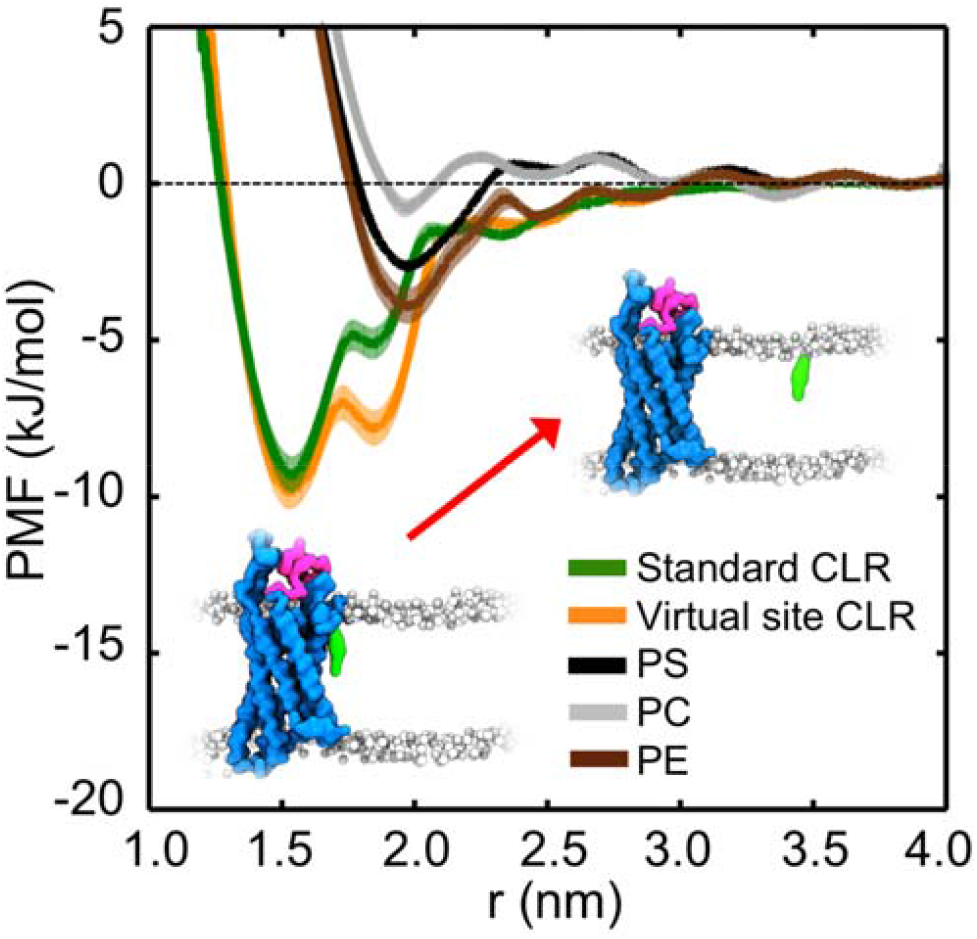
Potential of mean force calculations for lateral lipid interaction. PMF profiles for cholesterol interactions at the TM2/3e site. A profile obtained using the standard Martini cholesterol parameters (green) is compared to a profile obtained using the new virtual model (orange); and to the phospholipids PS (black), PC (grey), and PE (brown). Insets depict simulation snapshots of bound and unbound cholesterol. The shaded areas behind each curve indicates the standard deviation estimated from bootstrapping.

### SMO binds PIP_2_ lipids

A number of studies (e.g. (Dawaliby et al., 2015)) have suggested that phospholipids may allosterically regulate GPCRs. We therefore simulated SMO in a five-component lipid mixture mimicking an *in vivo* plasma membrane in composition and distribution (see previous sections for analysis of cholesterol interactions in this environment). These simulations revealed a high degree of interaction with negatively charged PIP2 lipids (Figure 7). This is of particular interest, as PIP2 has recently been shown to act as a positive allosteric modulator of Class A GPCRs, forming ‘encounter complexes’ enhancing G protein coupling by simultaneously contacting both structures as a ‘bridge’ or ‘molecular glue’ (Yen et al., 2018). The interaction of PIP_2_ with SMO occurred at defined regions on the intracellular portion of the protein, with multi-valent interactions predominately mediated *via* binding of the tri-phosphorylated headgroups of PIP2 to clusters of basic protein side chains (R257, K344, K356, K539, R546, and R547). This observation is likely to prove particularly intriguing should the binding of SMO to intracellular partners, such as G proteins, be structurally rationalized in the future. Furthermore in an *in vivo* context, we note that SMO is enriched near the base of primary cilia, a zone which contains high levels of PIP2. Ciliary phosphoinositides have been shown to regulate Hedgehog (Hh) signaling. Mutations in a 5-position phosphatase (Inpp5e) lead to alterations in the distribution of ciliary PIP_2_ and cause Joubert’s syndrome, a human ciliopathy characterized by impaired Hh signaling and human birth defects (Bielas et al., 2009; Chávez et al., 2015; Garcia-Gonzalo et al., 2015; Nakatsu, 2015). The simulation-based observation of direct PIP2 binding to defined regions of SMO is therefore of especial interest.

**Figure 7.**
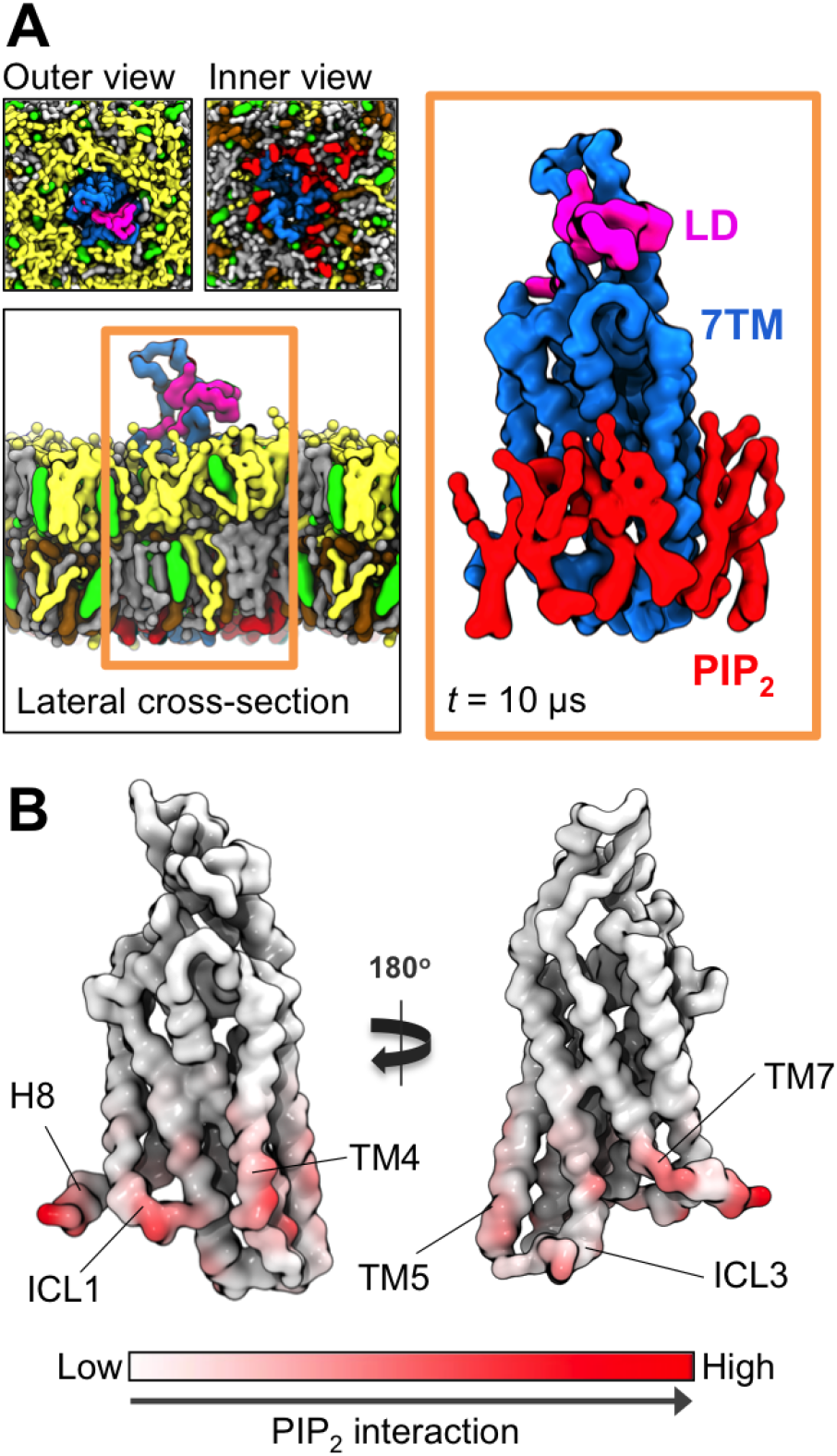
Plasma membrane mimetic simulations. (A) Final snapshots of SMO embedded in a PC (yellow), PE (grey), PS (ochre), PIP2 (red), cholesterol (green) membrane. Protein and lipids are rendered as surfaces. The formation of a PIP_2_-SMO complex (orange box) is apparent. (B) PIP_2_ contacts mapped onto structure, with each residue colored from white (no interaction), to red (high interaction). Contacts were calculated over an ensemble of 8 × 10 μs of CGMD, using a 6 Å distance cutoff. Each simulation was initiated from a different random distribution of lipids around the protein.

## Discussion

We observe direct interactions of cholesterol with the 7TMD of human SMO using molecular dynamics simulations at a range of resolutions, timescales, parameter sets, and membrane compositions. This is particularly intriguing given the recent discovery of a functional dependence of the truncated 7TMD of SMO on membrane cholesterol (Myers et al., 2017), as well as longstanding pathophysiological observations made in human developmental disorders (Blassberg et al., 2016; Cooper et al., 2003), and the emerging suggestion that statins may synergize with the
effects of vismodegib on SMO activity (Gordon et al., 2018). The identification of a well-defined cholesterol binding site located at TM2/3e is especially interesting. This cholesterol interaction is within the extracellular leaflet of the membrane. Recent structures of Ptch1 (Gong et al., 2018; Qi et al., 2018; Zhang et al., 2018) identify a potential interaction site between Patched1 (Ptch1) and cholesterol in a region that is also located in the extracellular leaflet. Whilst the mechanism of Ptch1 modulation of SMO remains uncertain, this suggests the view that Ptch1 may regulate SMO via extracellular leaflet cholesterol (Liu et al., 2017).

The cholesterol binding site at TM2/3e is distinct from the more transient interactions with the rest of the protein surface, as measured by density projections, contact analysis, analysis of occupancy dynamics, as well as free energy calculations. This site was consistently observed across multiple timescales, parameter sets, membrane compositions, and resolutions. In all cases cholesterol adopted a dynamic binding mode within this site, both undergoing frequent rapid exchange with bulk membrane cholesterol, and undergoing rotation about its long-axis whilst bound. This is consistent with the results of potential of mean force calculations at this site which yielded a well-depth of −10 kJ/mol. These observations point to a ‘greasy patch’ model for cholesterol binding to SMO, as observed for the Adenosine A2A receptor (A2aR) (Rouviere et al., 2017) and the dopamine transporter (Zeppelin et al., 2018).

In this context, it is useful to compare our estimates of the strength of SMO/cholesterol interactions with those obtained from other simulation studies of GPCRs, whilst bearing in mind the different methodologies and bilayer compositions employed in the various studies. Thus from analysis of extended (50 μs) equilibrium CG simulations of the β2AR and A2aR in a cholesterol-containing membrane, Geheden et al. (Genheden et al., 2017) estimated free energies of interaction of the order of −10 to −15 kJ/mol. From equilibrium (2 × 0.8 μs) atomistic MD simulations of the A2aR, Lee and Lyman (Lee and Lyman, 2012) estimated free energies of – 3 to −5 kJ/mol. A comparable analysis of our equilibrium CG simulations yielded a free energy of interaction of −6 kJ/mol. Given the detailed differences between the various studies, a conservative interpretation would be that cholesterol interactions with GPCRs are of the order of −5 to −10 kJ/mol, in contrast with estimated free energies of −10 to −40 kJ/mol for the interactions of a range of different membrane proteins with anionic phospholipids calculated by the same approach (see (Arnarez et al., 2016; Arnarez et al., 2013b; Domaήski et al., 2017; Domicevica et al., 2018; Gu et al., 2017; Hedger et al., 2016a; Hedger et al., 2016b) and Table S1).

A number of cholesterol interaction sites have been determined for Class A GPCRs (Sengupta and Chattopadhyay, 2015) the most well-established of which is perhaps the cholesterol consensus motif (CCM) suggested for the β2AR (Hanson et al., 2008). Our observations point to distinct cholesterol interaction patterns on SMO. This is perhaps not surprising. Although SMO, a Class F GPCR, bears a high degree of structural similarity to its Class A counterparts, it is distant in sequence with a sequence identity is < 10% (Wang et al., 2013). Additionally, an emerging pattern from the analysis of Class A GPCR cholesterol interactions is that these are often receptor specific, with limited sequence conservation (Gimpl, 2016). We did not observe any significant cholesterol interaction at the region of SMO corresponding to the CCM of the β_2_AR. This is perhaps unsurprising as the equivalent involved residues in SMO are differently distributed in space compared to those suggested for β_2_AR (Hanson et al., 2008), resulting in an arrangement for which it is difficult to explain how cholesterol could simultaneously contact the key residues of the motif.

Regarding cholesterol interaction sites more broadly, CRAC and CARC motifs have been suggested in some cases to form regions more conducive toward cholesterol binding (Epand, 2006; Fantini and Barrantes, 2013; Li and Papadopoulos, 1998). The general nature of the definition of these motifs results in their presence in most membrane proteins. Indeed, the simulated SMO construct contains a total of 13 of these motifs. However, most of these can be discounted as candidates simply because they are not membrane exposed, or their geometry is otherwise such that it is difficult to rationalize how cholesterol would bind. Comparison of the location of these motifs to the degree of per residue cholesterol contacts extracted from the CGMD simulations (Figure S10), revealed a degree of co-localization with the intracellular ends of TM4 and TM5. These regions exhibit lower levels of cholesterol occupancy and do not yield well-defined cholesterol density compared to TM2/3e. Nonetheless direct interaction is observed and the co-localization with the CRAC/CARC motifs is interesting.

Regarding lipid interactions in more complex multi-component membranes, the reproducibility of cholesterol interaction patterns in a plasma membrane mimetic is especially encouraging. This reproducibility suggests in this case, the absence of competition effects with other major plasma membrane lipids, as also supported by the PMF calculations. The absence of such effects is perhaps intuitively more likely for cholesterol, which is disparate in structure, physiochemical properties, and membrane insertion depth compared to its phospholipid counterparts and may thus be expected to interact with embedded proteins via rather different modes. The additional observation of the formation of a PIP2-SMO encounter complex in these membranes, similar to simulation/native mass spectrometry based observations for Class A GPCRs (Yen et al., 2018), is also intriguing and raises the prospect of potential involvement of PIP_2_ in modulation of SMO binding to putative intracellular interaction partners.

These results provide a testable hypothesis as to the manner of cholesterol interaction with SMO. We propose a number of routes for experimental testing of our observations. In the first instance, these predictions could be tested by site-directed mutagenesis coupled to subsequent functional assays. Such functional assays could either be conducted using signaling assays in cells (Luchetti et al., 2016), or indeed in minimal *in vitro* reconstituted nanodisc systems, where the activity of the truncated 7TMD has been shown to depend on cholesterol (Myers et al., 2017). Secondly, native mass spectrometry has shown recent tremendous utility in probing the specific binding of lipids to membrane proteins (Gupta et al., 2018; Laganowsky et al., 2014). Coupling this approach to a mutagenesis strategy could provide an exciting route to identify putative cholesterol interaction regions. A note of caution however, that some uncertainty remains as to the ability of native mass spectrometry to identify weak binding lipid species, including cholesterol. Most cases to date have focused on rigid high energy binding of species such as PIP (Laganowsky et al., 2014), PE (Patrick et al., 2018), and cardiolipin (Gupta et al., 2017), which may be expected to better survive detergent solubilization. This potential caveat is true also for crystallographic methods (Yeagle, 2014). In both of these cases it is worth highlighting also that mutagenesis of GPCRs to identify lipid binding is a non-trivial undertaking. One must be cognizant of the possible need for simultaneous mutation of clusters of binding site residues in order to evoke sufficient perturbation of the lipid binding site and preventing competition effects from neighboring residues (Yen et al., 2018), whilst at the same time remaining cognizant to the sensitivity of GPCR expression and folding to introduced perturbations. Careful choice of mutants and extensive controls are likely necessary. Thirdly, an exciting approach to identify cholesterol interaction sites is photo-sensitive chemical crosslinking, also referred to as ‘click’ assays (Hulce et al., 2013), which utilize photo-reactive cholesterol analogues to trap the interaction before subsequent mass spectrometry analysis. It is important to consider that whilst 1) tests whether the identified regions influence SMO signaling activity, 2) and 3) focus on testing simply whether cholesterol binds at particular sites, or not. It is possible of course that cholesterol interaction sites could have a range of biological functions besides influencing SMO signaling activity, including e.g. modulating lateral interactions with other biomolecules as has been seen for other lipid-protein interactions, and effects on stability. Both of which have been observed for other GPCRs (Prasanna et al., 2014; Zocher et al., 2012)

## Limitations

Accurate interpretation of our predictions requires a discussion of the limitations of the approaches used and currently available experimental data on which to base our model. The Martini model involves an inherent simplification of chemical detail (Marrink et al., 2007). This is a tradeoff made to access the time and length scales required for sufficient sampling of lipid-protein configurational space. Obtaining converged calculations of this nature in all-atom detail remains challenging without the use of specialized bespoke supercomputing resources (Shaw et al., 2008), and alternative enhanced sampling approaches (Domański et al., 2018). Importantly, we consider only one conformational state of the protein, for which structures have been determined. How might the location of cholesterol interaction sites mechanistically affect function? This is a challenging question to address at this time. In Class A GPCRs the major conformational transition between inactive and active occurs at the TM5, ICL3, TM6 interface (Dror et al., 2011). However significant conformational changes are possible at other regions such as ICL2 in the κ-opioid receptor (Che et al., 2018). The extensibility of these observations to Class F GPCRs remains uncertain. Should alternative conformational states of SMO emerge either from further structures and/or long timescale all-atom MD, it would be extremely interesting to re-visit cholesterol interactions and assess any possible deviations in interaction pattern.

Despite these limitations, the approaches discussed have achieved significant success in identifying a range of lipid interaction sites on membrane proteins, controlled and tested against experimental data including mass spectrometry (Gupta et al., 2017; Liko et al., 2016; Yen et al., 2018), crystallographic (Arnarez et al., 2013a; Schmidt et al., 2013; Van Eerden et al., 2017; Zeppelin et al., 2018), NMR spectroscopy (Duncan et al., 2018; Hedger et al., 2016b), and mutational functional data (Hedger et al., 2016a; Hedger et al., 2015; Stansfeld et al., 2009).

## Conclusions

These data provide key molecular level detail on the location and modes of direct cholesterol interaction with the 7TMD domain of SMO, a Class F GPCR of significant pharmaceutical interest, with an emerging intricate functional relationship with cholesterol. Identification and molecular level characterization of these sites is a first step toward understanding the mechanistic implications, and possible routes to therapeutic intervention *via* the design of small molecule mimetics, or the targeted control of cholesterol metabolism (Gordon et al., 2018).

## METHODS

### SMO Model Building

The SMO model used in simulations was based on the near full-length structure (PDB entry 5L7D) (Byrne et al., 2016). The primary goal of these simulations was to characterize cholesterol interactions with the transmembrane domain. As such the structure was truncated at position 191, yielding a construct (residues 192-549) consisting of the LD and 7TMD. This has previously been shown to be a stable unit for which multiple structures exist (Wang et al., 2014; Wang et al., 2013; Weierstall et al., 2014), and functionally viable in a membrane environment (Myers et al., 2017). Simulating only the LD and 7TMD construct enabled us to create smaller simulation boxes and expedite data collection. Side chain ionization states were modelled using pdb2gmx (Histidine) and PropKa (All other residues) (Olsson et al., 2011; Sondergaard et al., 2011). The N and C-termini were treated as neutral. The stabilizing and inactivating V329F mutation was left untouched. Intracellular loop 3 (occupied by the BRIL fusion in the 5L7D crystal structure) was modelled using coordinates from the PDB entry 4N4W (Wang et al., 2014). The protein structure was then energy minimized using the steepest descent algorithm implemented in GROMACS (Hess et al., 2008).

### Coarse-Grained Simulations

The minimized protein structure was converted to a CG representation using the Martini 2.2 force field (de Jong et al., 2013b; Monticelli et al., 2008). Tertiary structure was modelled using an Elnedyn network with a cutoff distance of 0.9 nm and a force constant of 500 kJ/mol/nm^2^. This approach prevents significant conformational deviations from the initial reference coordinates, while preserving local dynamics (Periole et al., 2009). The CG protein was centered in a simulation box of dimensions 10 × 10 × 13 nm, containing 280 randomly oriented 1-palmitoyl-2-oleoyl-sn-glycero-3-phosphocholine (POPC) lipids. The system was solvated using the standard Martini water model (Marrink et al., 2007), and neutralized with 0.15 M NaCl, before being subjected to 100 ns of CG simulation to permit the self-assembly of a lipid bilayer. This approach allows the protein to dynamically adopt its optimum orientation within the bilayer (Scott et al., 2008). Randomly selected POPC lipids were subsequently exchanged (Koldsø et al., 2014) for cholesterol molecules, to create mixed membranes of specified lipid composition (Table 1). Exchanges were only allowed outside a 2.5 nm cutoff distance from the protein surface, to avoid potential bias arising from fortuitous preplacement. This process was repeated for each individual repeat simulation, so as to create different random initial lipid configurations. An analogous process was performed for the 5-component plasma membrane mimetic simulations. Lipid compositions were chosen based on experimental lipidomics (van Meer et al., 2008). The standard Martini cholesterol parameters correspond to those of (Marrink et al., 2008), whilst the virtual site Martini cholesterol parameters were taken from (Melo et al., 2015). PIP2 parameters were created in-house as previously described (Stansfeld et al., 2009).

Temperature and semi-isotropic pressure were controlled at 310 K and 1 bar using the Berendsen barostat and Berendsen thermostat, with a coupling constants of 4 ps (Berendsen et al., 1984). Simulations of the CRD employed isotropic pressure coupling. Van der Waals interactions were smoothly shifted off between 0.9 nm and 1.2 nm. Modelling of electrostatics utilized the reaction field approach (Tironi et al., 1995), with a Coulomb cutoff of 1.2 nm and a potential shift modifier. Equations of motion were integrated with a 20 fs timestep, using the leapfrog algorithm implemented in GROMACS. Covalent bonds were constrained to their equilibrium values using the LINCS algorithm (Hess et al., 1997).

Simulation data was analysed using VMD (Humphrey et al., 1996), tools implemented in GROMACS (Hess et al., 2008), and in-house protocols. Protein-lipid contact analysis employed a cutoff distance of 0.6 nm, based on radial distribution functions for CG lipid molecules (Hedger et al., 2015). Likewise, annular lipids were calculated as those within 0.6 nm of the protein surface.

### Potential of Mean Force Calculations

Potential of mean force (PMF) calculations were performed using a protocol previously described (Hedger et al., 2016a; Hedger et al., 2016b). All 7TMD PMF calculations were conducted in a PC only bilayer with a single cholesterol molecule initially bound to the SMO TMD. This simple system was chosen to accelerate convergence during the PMF simulations (Domański et al., 2017). A one-dimensional reaction coordinate was generated using a steered molecular dynamics (SMD) simulation to pull the cholesterol from the bound to unbound state. Each cholesterol molecule was pulled away from the protein over a distance of 3.5 nm along a coordinate orthogonal to the protein surface. The SMD was performed at a rate of 0.1 nm/ns *(F_c_* = 1000 kJ/mol/nm^2^) *via* application of a force to the ROH particle of cholesterol. Within 7TMD PMF calculations, position restraints (F_c_ = 400 kJ/mol/nm^2^) were also applied in the *X-Y* plane to the backbone particles of V240, P369, and L464. These residues are distal from the respective cholesterol binding sites. In addition, weaker positional restraints (Fc = 50 kJ/mol/nm^2^) were applied to the ROH particle of cholesterol in the *Y* direction. Application of such restraints acted to prevent rotation of the protein, and translational “following” of cholesterol molecules as they were pulled away. Umbrella sampling simulations employed a window separation of 0.1 nm, using initial conformations extracted from the SMD simulation. Each window was run for 1 μs, with umbrella biasing potentials (F_c_ = 1000 kJ/mol/nm^2^) applied between the center of mass of the triad of restrained residues and the ROH particle of cholesterol. The subject lipid was treated separately from bulk lipids for temperature and pressure coupling. Approximately 32 umbrella sampling simulations were run per PMF. PMF profiles were constructed using the GROMACS implementation (g_wham) of the weighted histogram analysis method (WHAM) (Hub et al., 2010). Bayesian bootstrapping with 200 bootstraps was used to estimate the errors for each profile. Convergence was assessed by comparing profiles calculated from independent 100 ns segments of simulation time (Figure S1).

### Atomistic Simulations

Atomistic simulations were run using the GROMOS53a6 force field (Oostenbrink et al., 2004). The system was solvated using the SPC water model (van der Spoel et al., 1998) and neutralized with NaCl to a concentration of 0.15 M. Periodic boundary conditions were applied, with a simulation time step of 2 femtoseconds. A V-rescale thermostat (Bussi et al., 2007) was used to maintain temperature around 310 K, with a coupling constant of 0.1 ps, whilst pressure was controlled at 1 bar through coupling to a Parrinello-Rahman barostat (Parrinello and Rahman, 1981), with a coupling constant of 1 ps. Particle Mesh Ewald (PME) (Essmann et al., 1995) was applied to model long-range electrostatics. Van der Waals interactions were cut off at 1.2 nm. The LINCS algorithm was used to constrain covalent bond lengths (Hess et al., 1997).

## Author Contributions

G.H. set up, performed, and analyzed the simulations. G.H, C.S., R. R., and M.S.P.S. designed the research project. H.K. and M.C contributed analysis tools. G.H. and M.S.P.S. wrote the manuscript with contributions from all authors.

## Acknowledgements

We thank S. L. Rouse (Imperial College London, UK), B. Schiøtt (Aarhus University, Denmark), J. Schnell (University of Oxford, UK), and members of the M. S. P. S. laboratory. G.H. acknowledges the Medical Research Council for PhD and postdoctoral funding. Research in the M.S.P.S. laboratory is supported by the Wellcome Trust, BBSRC, and EPSRC. This project made use of time on ARCHER granted via the UK High-End Computing Consortium for Biomolecular Simulation supported by EPSRC (grant no. EP/L000253/1). CS is funded by Cancer Research UK (C20724/A14414) and the European Research Council (647278). RR is supported by grants GM106078 and GM118082 from the National Institutes of Health.

## Supporting Material

Supporting data figures and two movies are provided.

